# Selective alteration to CD4 T cell differentiation by heterozygous IRF4^L116R^ protects against neuroinflammation

**DOI:** 10.1101/2025.06.25.661400

**Authors:** Rebecca A. Jaeger, Nadia A. Roberts, Cynthia M. Turnbull, Sandali Seneviratne, Rebecca L. Buckland, Dominik Spensberger, Jessica A. Pettitt, Fiona D. Ballard, Anne Brüstle, Anselm Enders

## Abstract

The decision of naïve T cells to differentiate into a specific Th cell subset after antigen encounter is a critical pivot point of the immune response and, when out of balance, can lead to autoimmunity, allergy or immunodeficiency. Th subset differentiation is determined by the expression of transcription factors including IRF4. We here describe a point mutation (IRF4^L116R^) in the DNA-binding domain of IRF4 leading to a dysregulation of CD4-T cell subsets and their functions. This point mutation does not alter overall protein expression. In sharp contrast to IRF4-null mice, neither the CD4/CD8 ratio nor T cell activation and memory is altered in naïve mice carrying the point mutation. However, IRF4^L116R^ T cells are reduced in their capability to differentiate into Th1, Th17 and Treg cells in a dose-dependent manner, contrasting with the findings in IRF4^KO^ T cells. Particularly striking is the loss of Th1-differentiation in T cells from homozygous I*rf4*^*L116R/L116R*^ mice while T cells completely lacking IRF4 show no reduction in Th1 differentiation. Furthermore, despite maintained ability to generate Th17 cells, expression of the IRF4^L116R^ variant protects mice against disease development in the Th17 cell-dependent EAE mouse model of neuroinflammation. This contrasts with IRF4 knock-out mice, where expression of one wild-type allele of IRF4 is sufficient for full disease development. Together our results show that the L116R point mutation in IRF4 can selectively alter the differentiation and function of some CD4 T cell subsets and suggest that the L116R mutation is not a classical loss- or gain-of-function variant.

## INTRODUCTION

Interferon regulatory factor 4 (IRF4) is a key transcription factor (TF) expressed within the immune system, where it is critical for the development, differentiation, and function of various immune cell types(1). In T cells, IRF4 is particularly required for the differentiation of CD4^+^ T cells into T helper (Th) cell subsets and for their associated function in directing the immune system for targeted responses. IRF4 is indispensable for the differentiation of the inflammatory Th17 cells leading to a complete resistance to the experimental autoimmune encephalomyelitis model (EAE) of neuroinflammation in *Irf4*^*-/-*^ mice(2). IRF4 has also been reported to affect Th1, Th2, Th9, Tfh and Treg cells to differing extents (2-9). This broad role is due to the involvement of IRF4 at multiple points during differentiation, including in the regulation of lineage-specific TFs that determine cell fate or the expression of cytokines, both of which are necessary for the differentiation and function of CD4^+^ T cell subsets.

At the molecular level, IRF4 interacts with DNA through its DNA binding domain (DBD) and recognises the consensus 5’-GAAA-3’ sequence within IRF4-DNA binding motifs(10). By itself, as a homodimer, IRF4 has only weak DNA binding capabilities but can bind to motifs such as the IFN-stimulated response elements (ISREs) when highly expressed(11, 12). However, IRF4 predominantly binds DNA as a heterodimer together with binding partners such as PU.1(13) or BATF/JUN(14), amongst others. Heterodimerisation with these TFs allows higher affinity binding to DNA motifs such as erythroblast transformation-specific IRF composite elements (EICEs) or activating protein 1 IRF4 composite elements (AICEs), respectively(11, 15). Of note, IRF4 contains a C-terminal autoinhibitory domain. Upon the binding of IRF4 to PU.1, the interaction between the DBD and this autoinhibitory region is released, allowing the IRF4-PU.1 complex to bind DNA(16, 17). The affinity of IRF4 for specific DNA binding motifs, coupled with the varying presence of interacting TFs, allows for IRF4’s dynamic regulation of different functions across different cell types.

Point mutations in the DBD of IRF4 are frequently found in human cancers. One such mutation is the L116R mutation, changing the hydrophobic leucine (L) to a positively charged arginine (R). The location of L116 is predicted to be essential for heterodimer formation in the IRF4-PU.1 complex(18, 19). In addition to potentially altering the formation of heterodimers, the mutation has been shown to increase the IRF4 homodimer binding affinity 2-4-fold for common IRF4-DNA binding motifs such as ISRE, AICE, and EICE motifs(20). Recently, it was published that the overexpression of IRF4^L116R^ in purified naïve CD4^+^ T cells from a *Irf4*^*-/-*^ mouse caused an increase in IL-9 production, but a decrease in IL-13 expression after CD4^+^ T cells were directed to differentiate into the Th2 and Th9 lineages *in vitro*(21). This suggest that this point mutation may alter the requirement of IRF4 in T cell differentiation in a subset-specific manner. To allow the effect of IRF4^L116R^ on T cell subsets to be studied in a more physiological context as well as to enable *in vivo* functional assays, we have generated mice heterozygous and homozygous for this L116R mutation.

We here show that the L116R point mutation in the IRF4 DBD skews the B-T lymphocyte ratio, while CD4^+^ and CD8^+^ T cell populations remain unaltered in the periphery and thymus. However, the intrinsic capacity of T cells to differentiate into Th subsets is significantly affected by the point mutation in a dose-dependent manner. Furthermore, mice carrying this mutation on one allele are highly protected against the development of neuroinflammation in the EAE model despite normal *in vivo* generation of inflammatory T cell populations.

## RESULTS

### Normal expression of IRF4^L116R^

To confirm the expression of IRF4, we performed a Western blot using splenocytes isolated from *Irf4*^*+/+*^, *Irf4*^*L116R/+*^ and *Irf4*^*L116R/L116R*^ mice. This showed a normal level of IRF4 expression in all genotypes (Figure 1A) and as expected, an absence of IRF4 in cells from an *Irf4*^*-/-*^ mouse.

**Figure 1.**
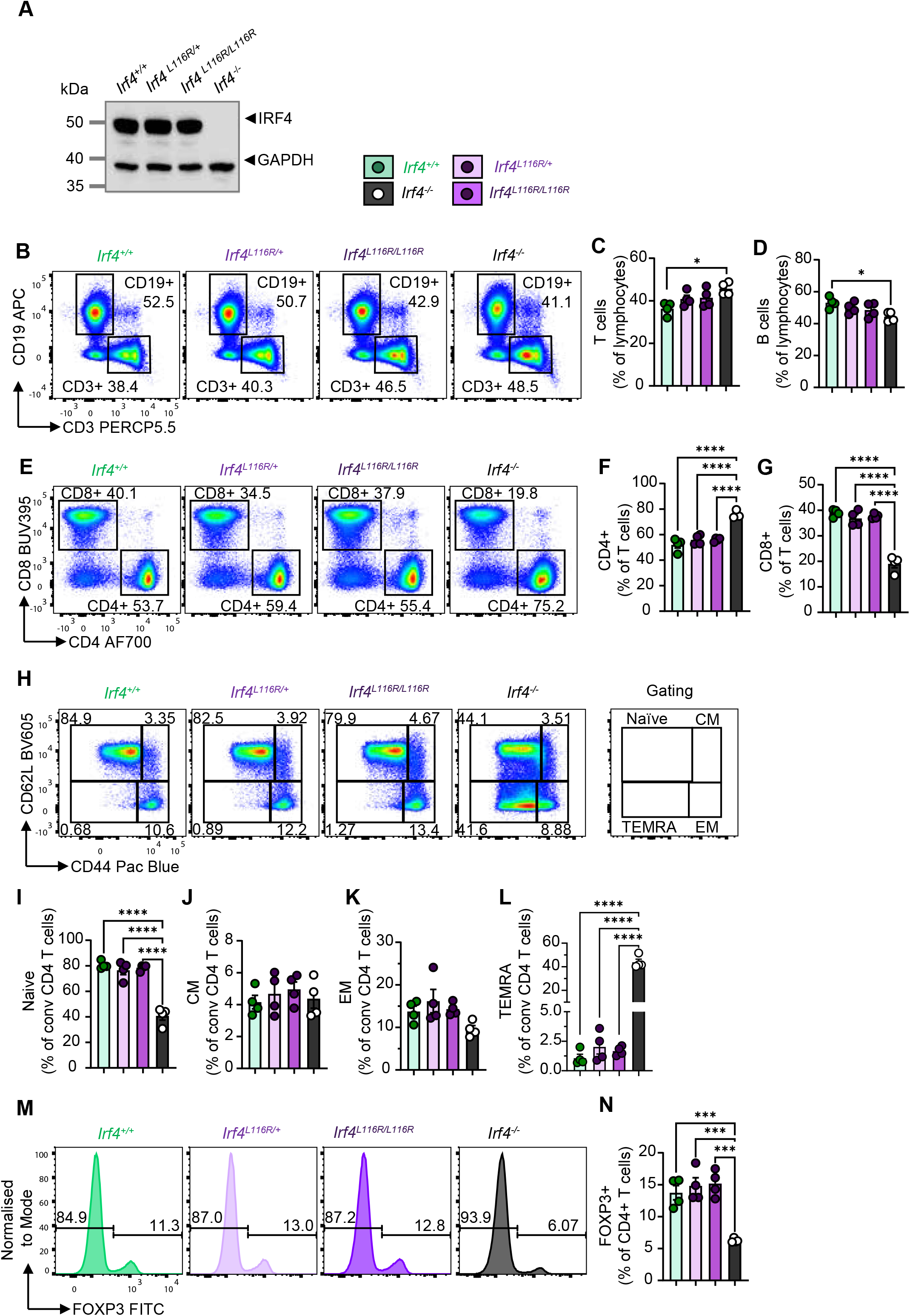
L116R IRF4 mutant expression and effect on splenic T lymphocyte proportions and activation *in vivo*. **(A)** Western blot of analysis showing IRF4 expression in splenocytes from *Irf4*^*+/+*^, *Irf4*^*L116R/+*^, *Irf4*^*L116R/L116R*^ and *Irf4*^*-/-*^ mice. **(B)** Representative flow cytometric profiles of CD3 vs CD19 expression in splenocytes for indicated genotypes. **(C-D)** Quantification of T (CD3+) and B (CD19+) cells in the spleen. **(E)** Representative flow cytometric plots of CD4+ and CD8+ T cells in the spleens for *Irf4*^*+/+*^, *Irf4*^*L116R/+*^, *Irf4*^*L116R/L116R*^ and *Irf4*^*-/-*^ mice. **(F-G)** Quantification of CD4+ and CD8+ cells as a proportion of CD3+ T cells in the spleen for indicated genotypes. **(H)** Representative flow cytometric profiles of spleen CD4+ naïve (CD62L^+^CD44^-^), central memory (CD62L^+^CD44^+^), effector memory (CD62L^-^ CD44^+^) and terminally differentiated effector memory (CD62L^-^CD44^-^) for *Irf4*^*+/+*^, *Irf4*^*L116R/+*,^ *Irf4*^*L116R/L116R*^ and Irf4^-/-^ mice. **(I-L)** Quantification of naïve, central memory (CM), effector memory (EM), and terminally differentiated effector memory (TEMRA) cells as a proportion of conventional (FOXP3-) CD4+ T cells in the spleen. **(M)** Representative histograms of FOXP3 expression in splenic CD4+ T cells for indicated genotypes. **(N)** Quantification of FOXP3+ Tregs as a proportion of CD4+ T cells in the spleen for indicated genotypes. Each dot is representative of a single mouse, n= 4. One-way ANOVA multiple comparisons adjusted with Tukey’s HSD are applied between groups.

### CD4^+^ T cell subsets

In line with results published from *Irf4*^*-/-*^mice, thymic T cell development was normal in heterozygous and homozygous *Irf4*^*L116R/+*^ and *Irf4*^*L116R/L116R*^ mice (Supplementary Figure 1). However, we found that homozygous *Irf4*^*L116R/L116R*^ mice trended towards an increased percentage of T cells/decreased percentage of B cells in spleen (Figure 1B-D) and peripheral blood (Supplementary Figure 2A-C), but this shift was not as pronounced as seen in *Irf4*^*-/-*^ mice. No change in the ratio of CD4^+^ and CD8^+^ T cells was observed in either spleen (Figure 1E-G) or blood (Suppl Fig 2D-F), whilst a shift towards CD4^+^ T cells was observed in the spleen of *Irf4*^*-/-*^ mice (Figure 1E-F) and, to a lesser extent, also in the blood (Suppl Fig 2 D-F). Furthermore, the absolute cell numbers of B and T cells in the spleen were unaffected (Supplementary Figure 3 A-D).

Analysis of T cell activation and memory subsets in naïve mice showed no major alterations in the frequency of CD4 memory cell populations either in the spleen (Figure 1 H-L) or blood (Suppl Figure 2 G-K) of heterozygous or homozygous *Irf4*^*L116R*^ mice. However, *Irf4*^*-/-*^ mice showed a marked increase in the proportions and absolute numbers of the CD44^-^CD62L^-^ TEMRA population in both the spleen and blood (Figure 1L, Supplementary Figure 2K and Supplementary Figure 3). Further, the frequency of Treg cells in *Irf4*^*L116R/+*^ and *Irf4*^*L116R/L116R*^ mice were comparable to *Irf4*^*+/+*^mice, contrasting with a significant reduction in Tregs seen in the spleen and blood of *Irf4*^*-/-*^ mice (Figure 1 M,N, Supplementary Figure 2 L,M), similar to previously published results(22). Together these results indicate that the L116R mutation acts as a hypomorphic loss-of-function mutation in the regulation of B- and T-cell numbers but has no effect on the development or homeostasis of regulatory T cells *in vivo*.

### In vitro CD4^+^ T cell differentiation

To investigate the T cell-intrinsic effect of IRF4^L116R^ on Treg differentiation, we performed *in vitro* CD4^+^ T cell differentiation analysis. *Irf4*^*L116R*^ T cells cultured under Treg-inducing conditions showed a slight dose-dependent reduction in Treg differentiation as measured by Foxp3^+^ cells on day 3 of culture (Figure 2A,B). This contrasts with the increased percentage of Treg cells in *Irf4*^*-/-*^ CD4 T cells as observed by us and as previously described by(2). The L116R mutation also introduced a gene dosage-dependent reduction in the formation of the pro-inflammatory Th1 cell subset, showing a specific effect not observed in *Irf4*^*-/-*^ CD4^+^ T cells (Figure 2C,D). *In vitro* differentiation into Th17 cells, another pro-inflammatory Th subset, was similarly reduced in a dose-dependent fashion in *Irf4*^*L116R*^ mice (Figure 2E, F). Nevertheless, the generation of Th17 cells was not entirely abolished in these mice, as opposed to what had been observed by us and others in *Irf4*^*-/-*^ mice (Figure 2E,F, and(2)).

**Figure 2.**
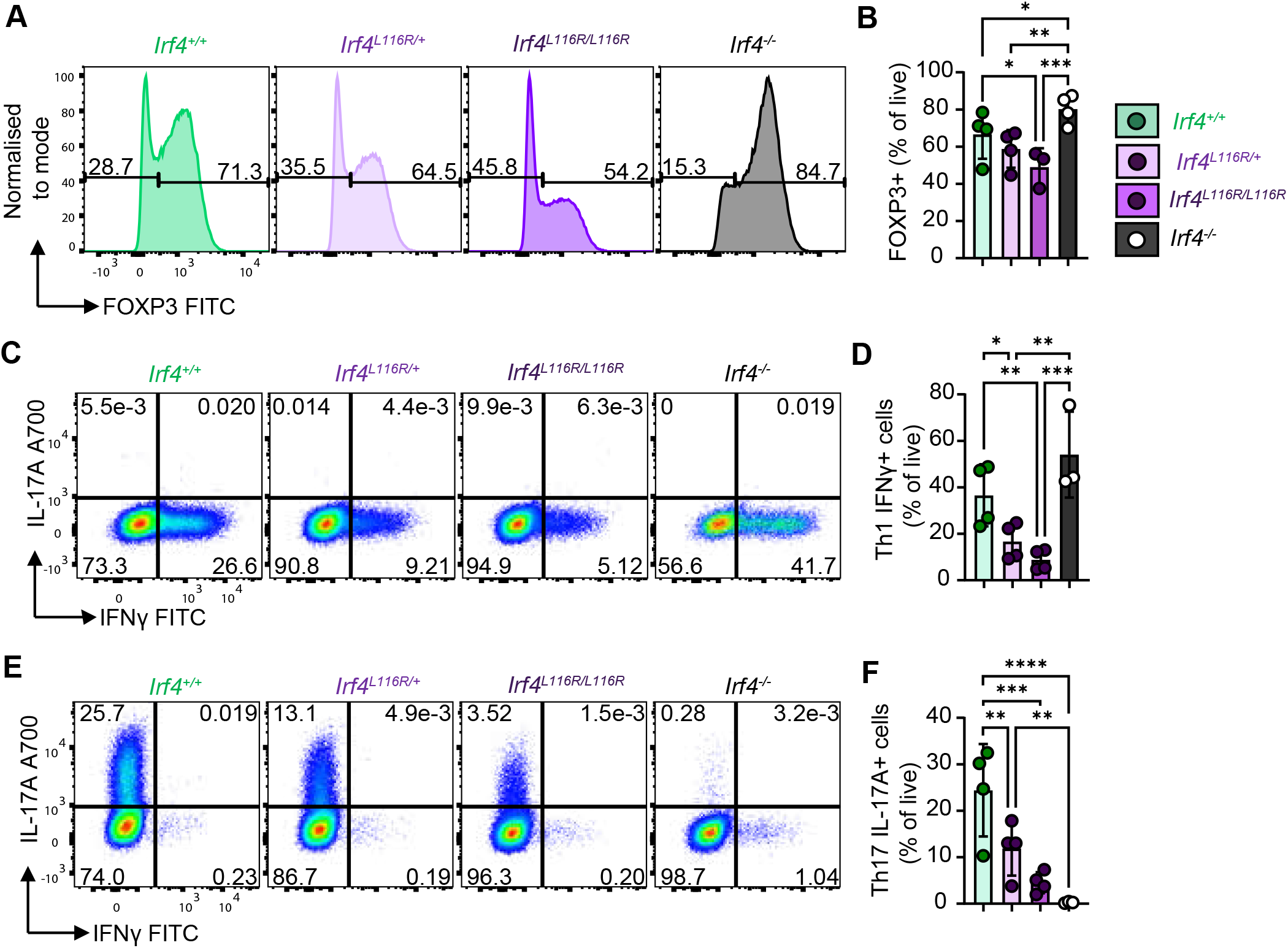
*In vitro* T helper cell differentiation with the L116R IRF4 mutation. **(A)** Representative histograms of FOXP3 expression in MACS sorted CD4+ Tnaive cells from *Irf4*^*+/+*^, *Irf4*^*L116R/+*^, *Irf4*^*L116R/L116R*^ and *Irf4*^*-/-*^mice under Treg differentiation stimuli. **(B)** Quantification of FOXP3+ cells following Treg differentiation for the indicated genotypes. **(C**,**E)** Representative flow cytometric plots of IL-17A and IFNγ expression in CD4+ Tnaive cells from *Irf4*^*+/+*^, *Irf4*^*L116R/+*^, *Irf4*^*L116R/L116R*^ and *Irf4*^*-/-*^mice under Th1 or Th17 differentiation stimuli, respectively. **(D**,**F)** Quantification of IFNγ and IL-17A expression. Each dot is representative of a single mouse, n=3-4. One-way ANOVA multiple comparisons are applied between groups using linear mixed models in R.

Together, these results show a suppression of regulatory T cell formation and a distinct effect on the differentiation of pro-inflammatory CD4^+^ T cell subsets by IRF4^L116R^.

### In vivo autoimmunity

To determine if these changes in T cell subsets have functional effects *in vivo*, we utilised the T cell-dependent EAE model(23). A complete protection against neuroinflammation in homozygous *Irf4*^*L116R/L116R*^ mice and a near-complete protection in heterozygous mice (Figure 3A,B) were observed. This is in sharp contrast to *Irf4*^*+/-*^ mice, which developed neuroinflammation and succumbed to EAE at a similar rate to *Irf4*^*+/+*^mice (Supplementary Figure 4A/B and(2)). T cell subsets in the blood at peak disease (day 21) revealed no differences in the activation status of the Th cells, as indicated by their expression of the T cell activation marker CD44 (Figure 3C,D) and their production of the cytokines IL-17A and IFNγ (Figure 3E-H). The unaltered levels of IL-17A in *Irf4*^*L116R*^ mice in EAE present an interesting contrast to the T cell differentiation results shown in Figure 2E,F, where IL-17A production by *Irf4*^*L116R*^ Th17 cells *in vitro* was drastically reduced. On the other hand, CD4^+^ T cells in *Irf4*^*-/-*^ mice did not upregulate IL-17 and EAE failed to develop in these mice (Supplementary Figure 4E,F). Of note, the production of GM-CSF, another cytokine associated with encephalitogenic Th cells(24), also remained unaffected in both heterozygous and homozygous *Irf4*^*L116R*^ mice (Figure 2I,J).

**Figure 3.**
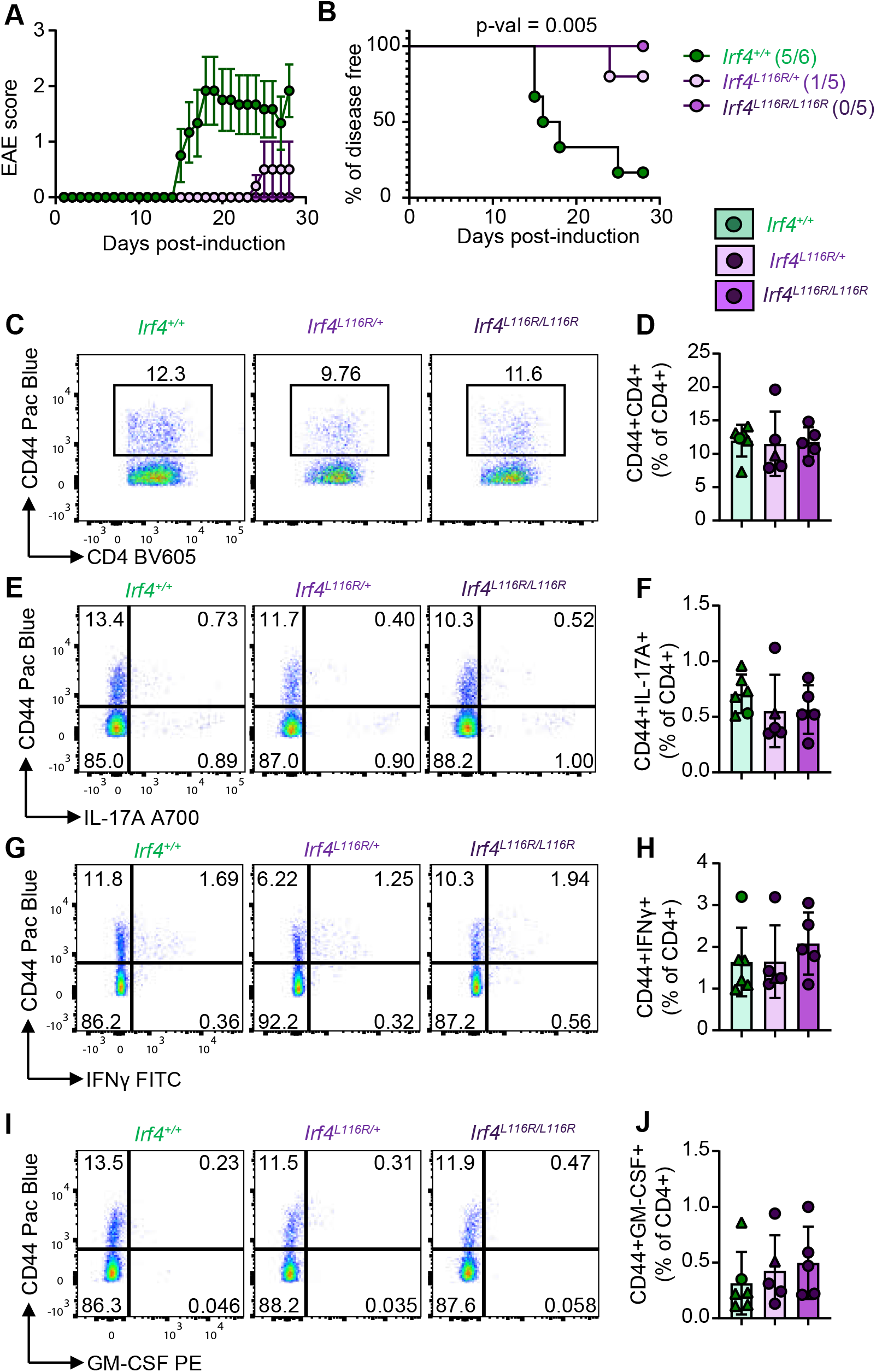
EAE with the L116R IRF4 mutation. **(A)** EAE score and **(B)** percentage of disease-free mice 28 days post-induction with MOG peptide for *Irf4*^*+/+*^, *Irf4*^*L116R/+*^, *Irf4*^*L116R/L116R*^ mice. Mice which become symptomatic are indicated in brackets. Representative flow cytometry plots for CD4 vs CD44 **(C)**, IL-17A vs CD44 **(E)**, IFNγ vs CD44 **(G)** and GM-CSF vs CD44 **(I)** expression in CD4+ T cells following PMA/ionomycin stimulation of blood samples collected on day 21 (EAE disease peak). Quantification of CD44+CD4+ **(D)**, CD44+IL-17A+ **(F)**, CD44+IFNγ+ **(H)** and CD44+GM-CSF+ **(J)** cells as a frequency of CD4+ T cells in the blood. Each dot is representative of a single mouse with symptomatic mice represented by triangles, n= 5-6 per genotype. Logrank (Mantel-Cox) test is applied for disease free survival curve comparison. One-way ANOVA multiple comparisons adjusted with Tukey’s HSD are applied between groups.

### Neuroinflammation

In line with the failure to induce EAE symptoms, we found a significant reduction in brain-infiltrating CD4^+^ T cells in *Irf4*^*L116R/L116R*^ mice. However, heterozygous mice displayed an infiltration of activated T cells despite being protected against EAE (Figure 4A,C). All T cells infiltrating the CNS in *Irf4*^*L116R/+*^ mice showed comparable levels of CD44 expression (Figure 4B) and in all genotypes infiltrating cells were able to produce IFNγ, IL-17A and GM-CSF (Figure 4D-I). Nevertheless, the proportion of cytokine producing CD4^+^ T cells was significantly reduced in the brains of *Irf4*^*L116R/L116R*^ mice. The percentage of Foxp3^+^ Treg cells was unaltered in all genotypes (Figure 4J, K).

**Figure 4.**
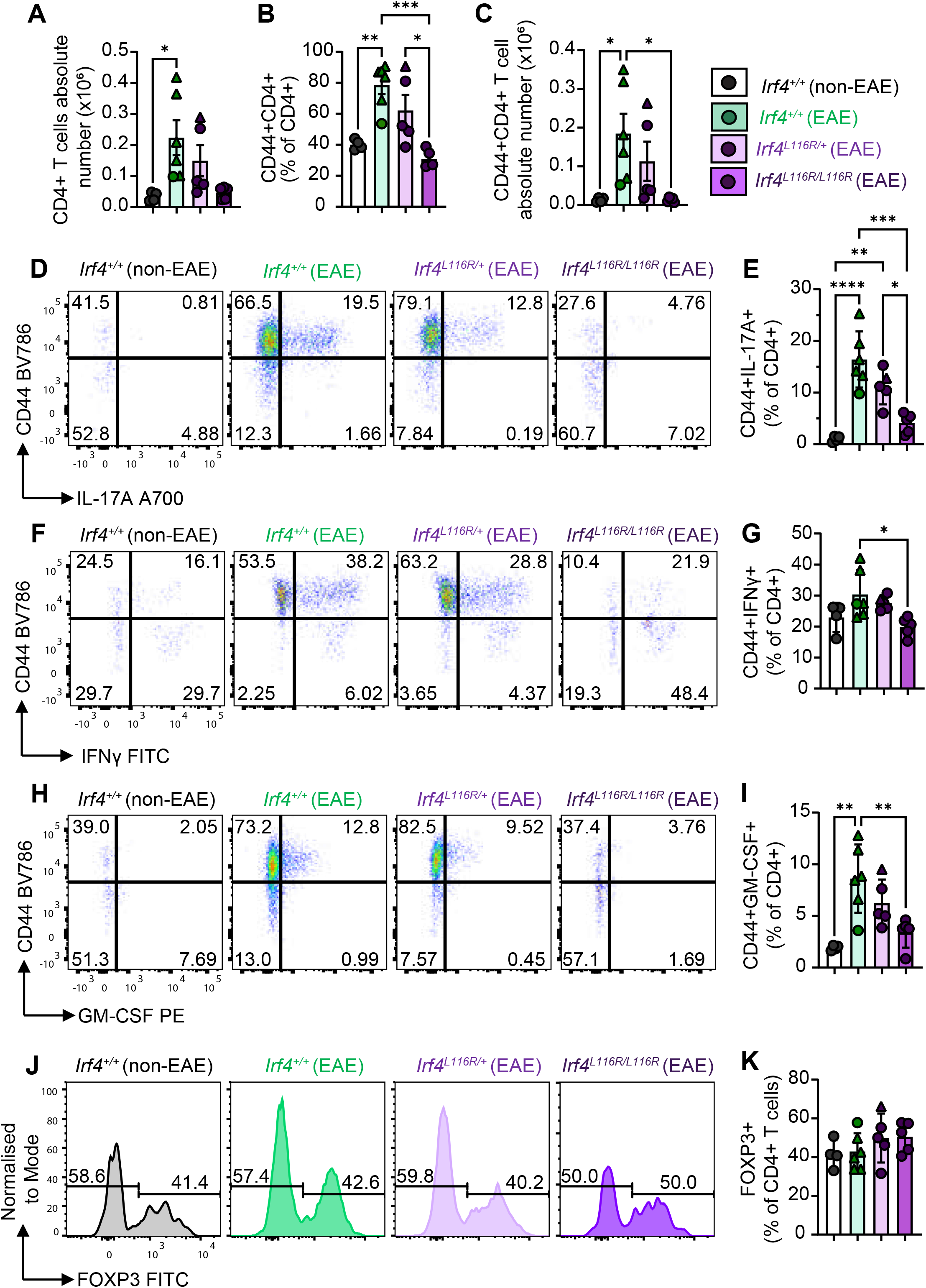
CNS infiltration during EAE with the L116R IRF4 mutation. **(A)** Absolute CD4+ T cell numbers, **(B)** CD44+CD4+ cell frequency of CD4+ T cells, and **(C)** absolute CD44+CD4+ T cell numbers in the CNS of *Irf4*^*+/+*^, *Irf4*^*L116R/+*^, *Irf4*^*L116R/L116R*^ mice 28 days post-induction with MOG peptide. Representative flow cytometry plots for IL-17A vs CD44 **(D)**, IFNγ vs CD44 **(F)** and GM-CSF vs CD44 **(H)** expression in CD4+ from PMA/Ionomycin stimulated CNS cells. Quantification of CD44+IL-17A+ **(E)**, CD44+IFNγ+ **(G)** and CD44+GM-CSF+ **(I)** cells as a frequency of CD4+ T cells in the CNS. Flow cytometric profiles **(J)** and quantification **(K)** of FOXP3 expression in CD4+ T cells in the CNS. Each dot is representative of a single mouse with symptomatic mice represented by triangles, n= 4-6 per genotype. One-way ANOVA multiple comparisons adjusted with Tukey’s HSD are applied between groups.

## Discussion

In contrast to an IRF4-null mutation, IRF4^L116R^ does not drastically affect the development of CD4^+^ and CD8^+^ T cells. However, we show that the L116R mutation in the DBD of IRF4 selectively alters the formation of specific Th subsets. Remarkably, the effect of the mutation is clearly different to a null allele. For example, we find that Th1 cells are strongly reduced, an effect not seen in the absence of IRF4 and T cells with either one or two copies of IRF4^L116R^ show some retained capacity to differentiate into Th17 cells, contrasting with the complete block in Th17 development in the absence of IRF4(2).

It had previously been shown that a single copy of wild-type IRF4 was sufficient to allow the development of EAE symptoms equivalent to *Irf4*^*+/+*^mice(2). In contrast, *Irf4*^*L116R/+*^ mice, which also retained one wild-type *Irf4* allele, were nearly completely protected against development of EAE symptoms, despite the maintained formation of pro-inflammatory Th17 cells *in vivo* following EAE induction. (18)

As such, the combination of effects on *in vitro* T cell subset differentiation and *in vivo* EAE development can neither be explained by a general loss of function effect of the mutation, nor by a dominant-negative effect. The L116R mutation has been shown to increase affinity to known DNA-binding motifs(20). However, the mutation does not appear to produce effects that are suggestive of a general gain of function in this study. The complex alterations observed in the differentiation and function of Th cell subsets suggest that the L116R mutation has a multimorphic effect similar to what has recently been described for two other mutations in the DBD of IRF4(25, 26). IRF4^L116^ has previously been identified as a crucial residue for interaction between IRF4 and PU.1(18), a key binding partner of IRF4 in some Th subsets and other immune cells. Because of this we modelled the effect of the L116R mutation on IRF4 and found that the Arginine reaches further across the DNA to PU.1 (Supplementary Figure 5). This suggests that the mutation not only alters binding to DNA but may also alter binding to PU.1.

Importantly, PU.1 has been shown to be essential in Th9 differentiation and Th9 cells have a known role in the induction of neuroinflammation in EAE(27). As such, alterations to heterocomplex formation with PU.1 may explain the observed protection against EAE symptoms in the *Irf4*^*L116R*^ mice. However, since both IRF4 and PU.1 are also co-expressed in other immune cells such as macrophages, it is difficult to exclude possible indirect effects like alterations to the balance of macrophage pro- and anti-inflammatory effector functions in the CNS, that might protect these mice against EAE(28).

However, though the precise molecular mechanism of the L116R mutation has yet to be elucidated, our data show for the first time a genetic separation of effects on Th subset differentiation and protection against neuroinflammation through a single point mutation in IRF4.

## EXPERIMENTAL PROCEDURES

### Mice

*Irf4*^*L116R*^ mice were generated in-house through CRISPR-Cas9-mediated genome editing. Mouse zygotes were obtained by mating superovulated C57BL/6NCrl females and WT C57BL/6NCrl males. The guide RNAs (gRNA1 5’ ATGGGTCAGAGATATCCAGC 3’ and gRNA2 5’ GGATATCTCTGACCCATACA 3’) were mixed with TrueCut Cas9 (Invitrogen) and 149bp ssODNs. The microinjection complex contained 100 ng/μl Cas9, 15 ng/μl of each gRNA, and 20 ng/μl ssODNA. The ribonucleoprotein complex mix was microinjected into the cytoplasm or pro-nuclei of zygotes, and the injected embryos were incubated overnight at 37°C at 5%CO2 in M16 medium (Merck) until they reached a two-cell stage. Two-stage embryos were transferred into pseudo-pregnant females the following day. Fifteen embryos were transferred unilaterally to each Swiss pseudo-pregnant recipient. All newborn pups were genotyped using specific primers (Fwd 5’ CCTTCCTTGCTGAAATGCAATC 3’ and Rev 5’ TCCAGATCCTAATTGAATGGAG 3’), and the amplicons were Sanger sequenced to confirm the correct ssODN integration. *Irf4*^*-/-*^ mice contain a 47 base pair deletion leading to a premature stop codon, with lack of protein expression confirmed through Western blot as previously described(25).

All mice were maintained on a C57BL/6NCrl background and housed in specific pathogen-free conditions at the Australian Phenomics Facility (Australian National University, ACT, Australia). Unless stated otherwise, mice were aged between 5 and 23 weeks, with both male and female mice used for experiments. All experiments were performed under ethics protocols 2020/45, 2021/49 and A2023/311 approved by the Australian National University Animal Ethics and Experimentation Committee.

### Experimental Autoimmune Encephalomyelitis

EAE was induced in 10-16-week-old *Irf4*^*L116R/+*^, *Irf4*^*L116R/L116R*^, *Irf4*^*+/-*^, *Irf4*^*-/-*^ and wild-type littermate control mice through subcutaneous immunisation with 115μg myelin oligodendrocyte glycoprotein 35-55 (MOG_35-55_, Peptides & Elephants) in emulsion with Complete Freund’s Adjuvant (Sigma-Aldrich) supplemented with 4mg/mL heat-killed *Mycobacterium tuberculosis* (BD). On days 0 and 2 of model, mice were intraperitoneally injected with 400ng Pertussis toxin (List Biological Laboratories). Clinical scoring of disease severity was performed daily on the following scale: 0 = no paralysis; 1 = flaccid tail and/or ataxia; 2 = hind limb weakness (loss of rolling reflex); 3 = hind limb paralysis; 4 = hind and front limb paralysis; 5 = moribund.

### Isolation of lymphocytes from tissue

For *ex vivo* flow cytometry analysis, spleens were prepared by passing through a 70μM filter and washing in PBS containing 2% FBS. Cells were then resuspended in 3mL RBC lysis buffer (150mM NH_4_Cl, 10mM NaHCO_3_, 1mM EDTA) for 3 minutes at room temperature and washed twice in PBS 2% FBS prior to antibody staining.

For isolation of CNS-infiltrating leukocytes during EAE, mice were perfused with 20mL PBS prior to collection of brain and spinal cord. CNS tissue was then mechanically dissociated and digested for 22 minutes at 37°C 5% CO_2_ in 50μg/mL collagenase P (Roche) and 10μg/mL DNAse I (Roche) in RPMI (Gibco). Cell digests were then passed through a 70μM filter, pelleted through centrifugation and layered through a Percoll (Sigma, 40% and 70% in DMEM) density gradient. Following collection of leukocytes from the interface, cells were washed twice in PBS containing 2% FBS prior to surface antibody staining, PMA and ionomycin restimulation and intracellular staining.

### T cell isolation and differentiation

Naïve CD4^+^CD62L^+^ cells were isolated from the spleens and lymph nodes of 8-12 week old *Irf4*^*L116R/+*^, *Irf4*^*L116R/L116R*^ and *Irf4*^*+/+*^ littermates as well as *Irf4*^*-/-*^ mice using magnetic-activated cell sorting (naïve CD4^+^ T cell isolation kit, Miltenyi Biotec) according to manufacturer’s instructions. For *in vitro* T cell differentiation, cells were cultured in a 96-well plate (150,000 cells per well) for 72 hours in the presence of 3μg/mL plate-bound αCD3 and 2μg/mL αCD28 (both BioXCell). Th1 cells were generated via the addition of 4ng/mL IL-12 (Miltenyi Biotec), 5ng/mL IL-2 (Peprotech) and 2.5μg/mL αIL-4 (BioXCell). Th17 cells were polarised using 30ng/ml IL-6 and 0.5ng/ml hTGFβ (both Miltenyi Biotec) in the presence of 2.5μg/ml αIL-4 and 10μg/ml αIFNγ (BioXCell), Treg cell polarisation conditions included 5ng/mL IL-2, 10μg/mL αIFNγ, 2ng/mL hTGFβ and 2.5μg/ml αIL-4.

Cells were cultured using complete IMDM (ThermoFisher, supplemented with 10% FBS, penicillin, streptomycin and glutamine) as basal media and incubated at 37°C 5% CO_2_.

### Restimulation and intracellular cytokine stain

For assessment of cytokine production directly *ex vivo*, cells were stained for surface markers for 30 minutes prior to restimulation with 50nM PMA (Sigma) and 500ng/mL ionomycin (Sigma) in the presence of 1:1000 volume Golgi stop (BD) for 4.5 hours at 37°C 5% CO_2_. Alternatively, *in vitro* differentiated Th cells were stimulated as above without an additional surface stain. Cells were then washed in PBS and incubated with Fixable Viability Dye ef780 (eBioscience, 1:1000) for 20 minutes at room temperature. Fixation, permeabilisation and intracellular staining were then performed using the FOXP3/transcription factor buffer kit (ThermoFisher) according to manufacturer’s instructions.

### Flow cytometry

Prior to staining for surface markers, cells were incubated for 10-15 minutes at room temperature in Fc block (BD, 1:100 in PBS 2% FBS). Antibody cocktail was then added, and cells incubated for a further 30 minutes at 4°C. The following antibodies were used for flow cytometry: CD19-APC (1D3, BD Biosciences), CD19-BUV395 (1D3, BD Biosciences), CD25-PE (PC61.5, eBioscience), CD3-PerCP CY5.5 (17A2, Biolegend), CD4-AF700 (RM4-5, BD Biosciences), CD4-BV605 (RM4-5, BD Pharmingen), CD44-PacificBlue (IM7, Biolegend), CD44-APC (IM7, BD Biosciences), CD44-BV786 (IM7, BD Biosciences), CD62L-BV605 (MEL-14, Biolegend), CD69-PE-Cy7 (H1.2F3, Biolegend),CD69-FITC (H1.2F3, Biolegend),CD8-BUV395 (53-6.7, BD Biosciences), CD8-PECy7 (53-6.7, Biolegend), FOXP3-FITC (FJK-16S, eBioscience), GM-CSF-PE (MP1-22E9, Biolegend), IFNγ-FITC (XMG1.2, BD Biosciences), IL-13-PECy7 (eBio13a, eBioscience), IL-17A-AF700 (TC11-18H10, Biolegend), IL-9-APC (RM9A4, Biolegend).

Cells were then washed in PBS 2% FBS and acquired using a BD LSRII or LSRFortessa flow cytometer at the Cytometry, Histology and Spatial Multiomics (CHASM) facility, ANU, with FACSDiva (BD Biosciences). Analysis was done using FlowJo v10 (BD Biosciences).

### Western blot

For IRF4 gene expression analysis western blotting was conducted on T cells from spleen and lymph nodes, activated for 16hrs in complete IMDM media with 2μg/mL αCD28 (BioXCell) and 5ng/mL IL-2 (Peprotech) on 3μg/mL αCD3 (BioXCell) coated plates. Cells were lysed in lysis buffer (20 mM Tris-HCL, pH 7.4, 150 mM NaCl, 1% Triton, 0.5% sodium deoxycholate, 0.1% sodium dodecyl sulfate (SDS)) supplemented with protease inhibitor (Roche) and sample loading dye buffer (50 mM Tris, pH 6.8, 100 mM dithiothreitol (DTT), 2% SDS, 0.1% bromophenol blue, 10% glycerol).

Proteins were separated on 4–12% polyacrylamide gradient Bolt Bis-Tris gel (Invitrogen) in MOPS running buffer (Invitrogen). Following semi-dry electrophoretic transfer of proteins onto polyvinyldifluoride (PVDF) membranes (IPVH00010, Millipore), membranes were blocked in 5% skim milk in tris-buffered saline with Tween-20 (TBST) and incubated overnight with primary antibodies against IRF4 (1:1000 dilution, 62834, Cell Signaling Technologies) and GAPDH (1:4000 dilution, ab8245, Abcam). PVDF membranes were then incubated with anti-rabbit (1:5000 dilution, 111035144, Jackson ImmunoResearch) or anti-mouse (1:5000 dilution, 115035146, Jackson ImmunoResearch) horseradish peroxidase-conjugated secondary antibodies for 1 hour and proteins were visualized using Clarity Western ECL Substrate (170-5061, BioRad) and the ChemiDoc Touch Imaging System (BioRad). Immunoblots were analyzed using ImageJ Software.

### Statistical analysis

Data was analysed using FlowJo10 software. Statistical analyses were performed using GraphPadPrism10 (single experiments) or RStudio (pooled experiments), as described. Unless stated otherwise, one-way ANOVAs were completed with multiple comparisons between groups (Tukey’s method). Linear mixed models applied in R for pooled experiments used experimental repeat number as the blocking factor. Disease free survival curves completed for EAE analyses were completed using a log-rank (Mantel-Cox) method.

## DATA AVAILABILITY

All data is contained within the manuscript. Mouse strains are available upon request (AE).

## ACKNOWLEDGMENT

We would like to acknowledge the Phenomics Translational Initiative for support of the Gene Targeting Facility (Rachael Milne, Harish Padmanabhan, and Neel Thakkar) at the Australian Phenomics Facility. We thank Dr Harpreet Vohra and Mick Devoy from the Cytometry, Histology and Spatial Multiomics (CHASM) Facility at the John Curtin School of Medical Research for cell sorting and FACS analysis. We thank Ms Yanran Fan for careful editing of the manuscript.

## Abbreviations

AICE: activating protein 1 IRF4 composite elements
BATF: basic leucine zipper transcription factor, ATF-like
CNS: central nervous system
DBD: DNA binding domain
EAE: experimental autoimmune encephalomyelitis
EICE: erythroblast transformation-specific IRF composite element
GM-CSF: granulocyte macrophage-colony stimulating factor
IFN: interferon
IL: interleukin
IRF4: interferon regulatory factor 4
ISRE: IFN-stimulated response element
JUN: jun proto-oncogene
KO: knockout
PU.1: Spi-1 proto-oncogene
TEMRA: T effector memory cells re-expressing CD45RA
TF: transcription factor
Tfh: T follicular helper
Th: T helper
Treg: T regulatory

This article contains supporting information

