## Supplemental Figures for "Selective alteration to CD4 T cell differentiation by heterozygous IRF4^L116R^ protects against neuroinflammation"

Supplementary figures-

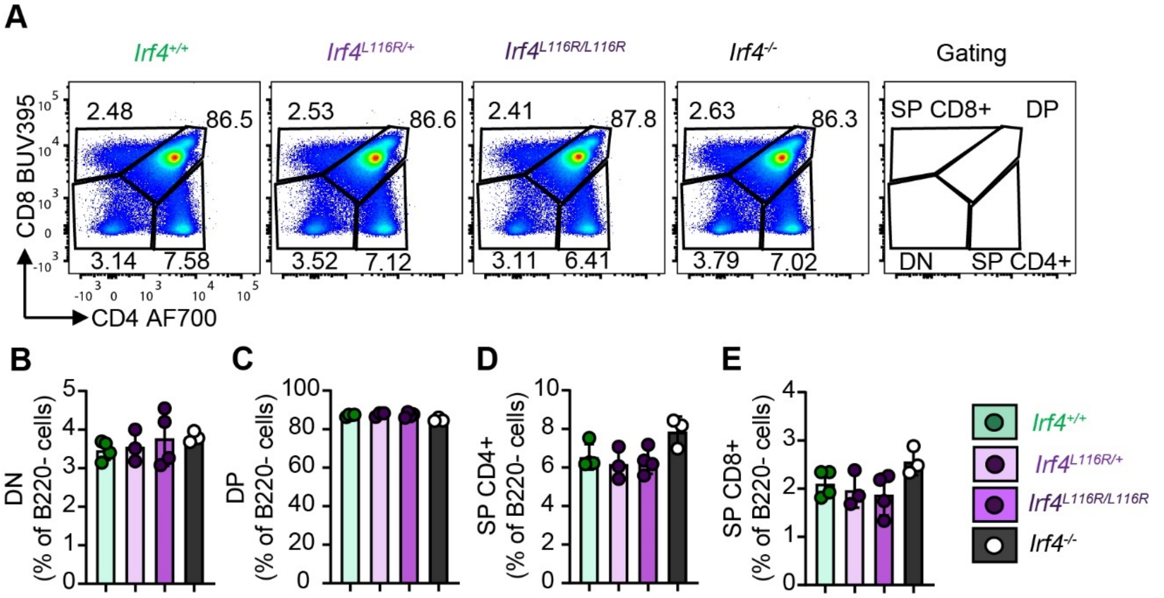


**Supplementary Figure 1-T cell development in the thymus with the L166R IRF4 mutation**
**(A)** Representative flow cytometric plots of CD8 vs CD4 expression for the indicated genotypes within the thymus. **(B-E)** Quantified frequencies of double negative (DN) **(B)**, double positive (DP) **(C)** and single positive (SP) CD4+ **(D)** and CD8+ **(E)** cells as a proportion of B220- cells in the thymus. Each dot is representative of a single mouse, n= 3-4. One-way ANOVA multiple comparisons are applied between groups.

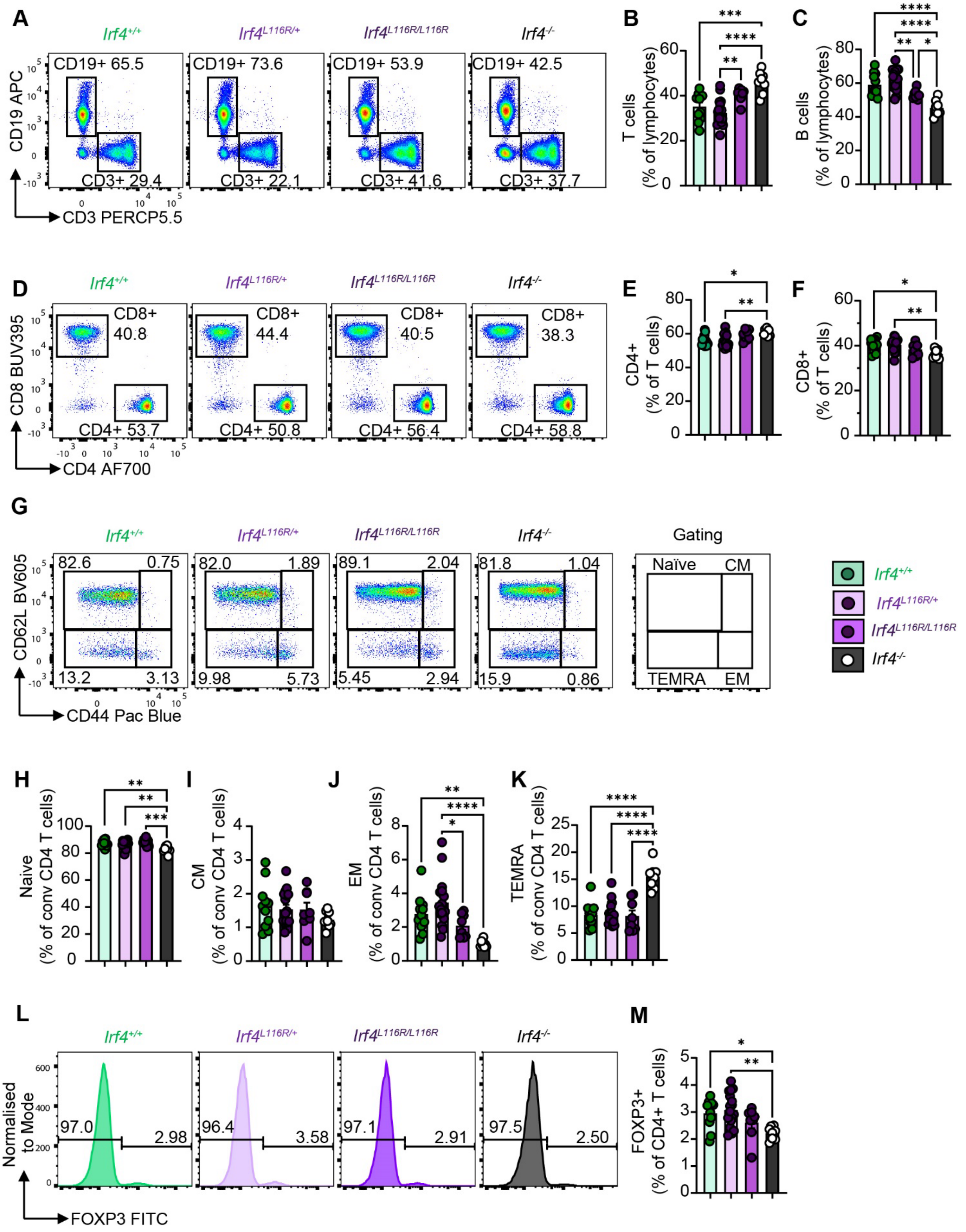


**Supplementary Figure 2 - L116R IRF4 mutant effect on T lymphocyte proportions and activation in blood *in vivo***

**(A)** Representative flow cytometric profiles of CD3 vs CD19 expression in blood for indicated genotypes. **(B-C)** Quantification of T (CD3+) and B (CD19+) cells in the blood for indicated genotypes. **(D)** Representative flow cytometric plots of CD4+ and CD8+ T cells in the blood of *Irf4*$^{+/+}$, *Irf4*$^{L116R/+}$, *Irf4*$^{L116R/L116R}$ and *Irf4*$^{-/-}$ mice. **(E-F)** Quantification of CD4+ and CD8+ cells as a proportion of CD3+ T cells in the blood for indicated genotypes. **(G)** Representative flow cytometric profiles of blood CD4+ naïve (CD62L$^{+}$CD44$^{-}$), central memory (CD62L$^{+}$CD44$^{+}$), effector memory (CD62L$^{-}$CD44$^{-}$) and terminally differentiated effector memory (CD62L$^{-}$CD44$^{-}$) for *Irf4*$^{+/+}$, *Irf4*$^{L116R/+,}$ *Irf4*$^{L116R/L116R}$ and Irf4$^{-/-}$ mice. **(H-K)** Quantification of naïve, central memory (CM), effector memory (EM), and terminally differentiated effector memory (TEMRA) cells as a proportion of conventional (FOXP3-) CD4+ T cells in the blood. **(L)** Representative histograms of FOXP3 expression in the blood for indicated genotypes. **(M)** Quantification of FOXP3+ Tregs as a proportion of CD4+ T cells in the blood for indicated genotypes. Each dot is representative of a single mouse, n= 8-17. One-way ANOVA multiple comparisons are applied between groups.

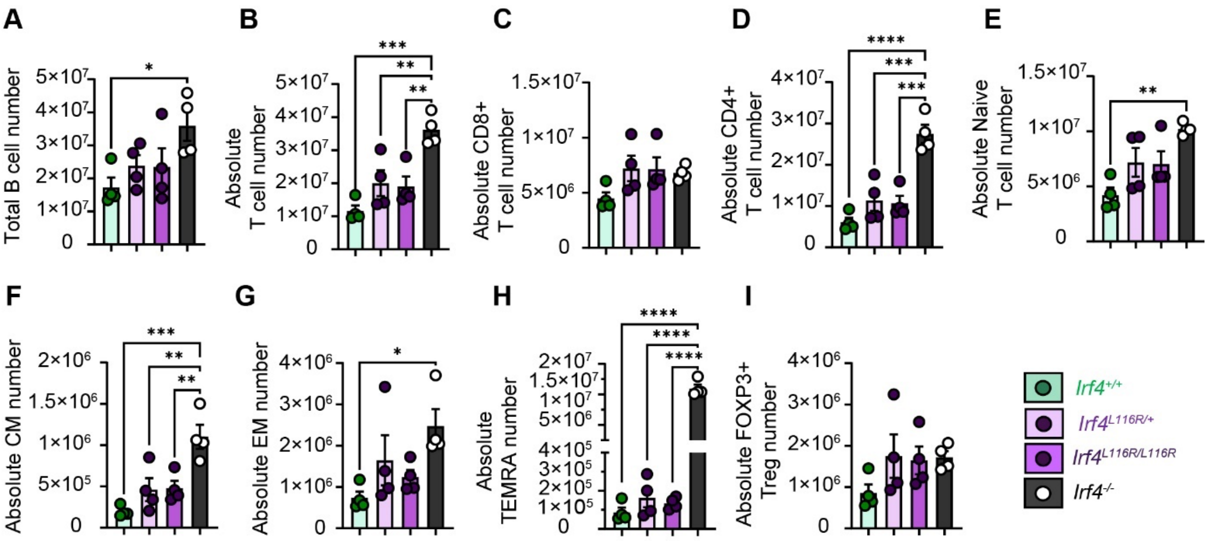


**Supplementary Figure 3-Absolute splenic cell numbers for T cell populations with the L116R IRF4 mutation**

**(A-E)** Absolute B cell, T cell, CD8+ T cell, CD4+ T cell and Naïve T cell numbers from splenic samples of the WT, Irf4$^{L116R/+,}$ Irf4$^{L116R/L116R}$ and Irf4$^{-/-}$ mice. **(F-I)** Absolute numbers for central memory (CM), effector memory (EM), terminally differentiated effector memory (TEMRA) and Tregs (FOXP3+) cells for the indicated genotypes. Each dot is representative of a single mouse, n=4. One-way ANOVA multiple comparisons are applied between groups.

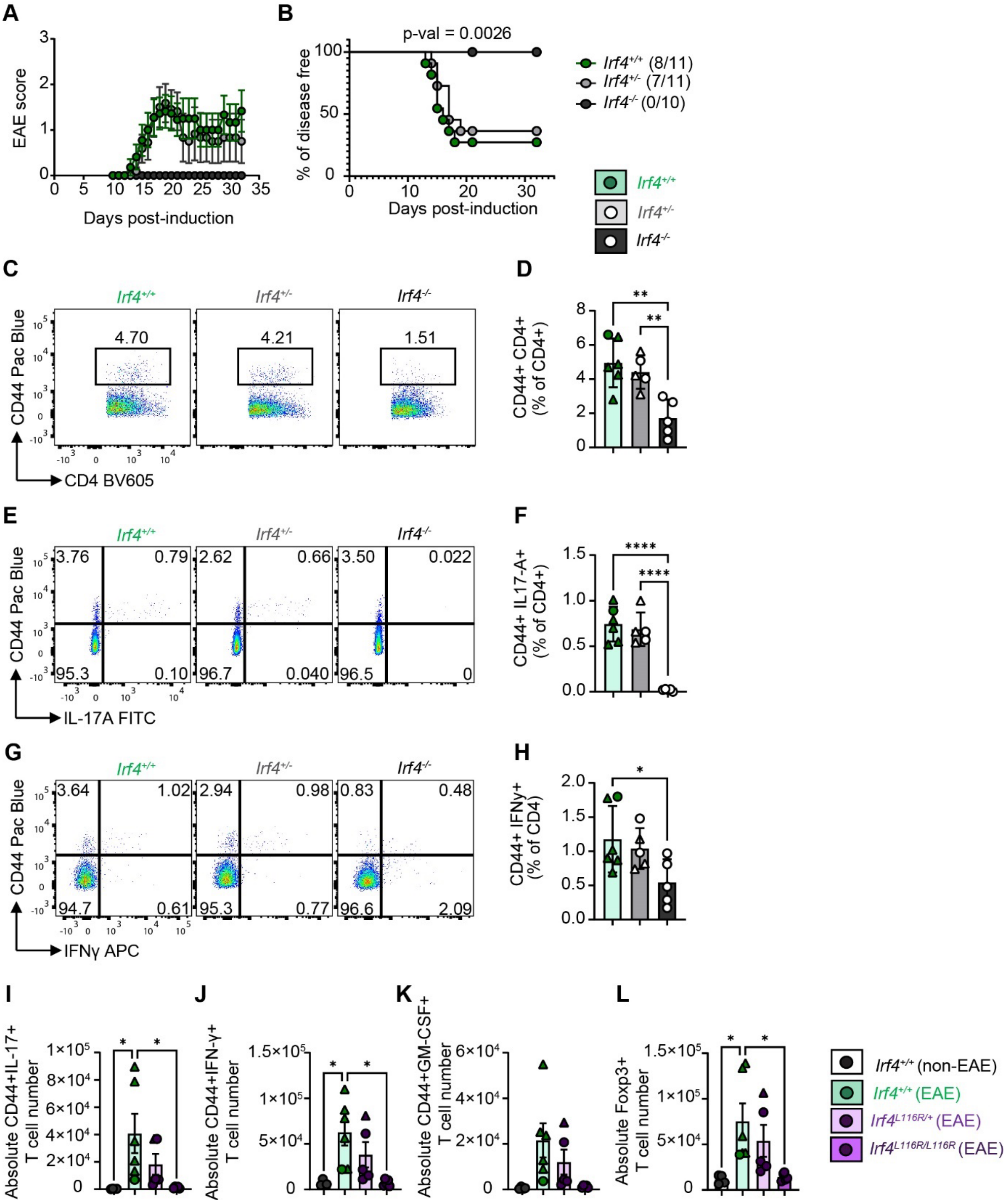


**Supplementary Figure 4- EAE with the IRF4 deletion and the absolute cell counts for CNS infiltration during EAE for L116R mutations.**

**(A)** EAE scores and **(B)** percentage of disease-free mice for *Irf4*$^{+/+}$, *Irf4*$^{+/-}$ and *Irf4*$^{-/-}$ mice following induction for EAE with MOG peptide. Mice which become symptomatic are indicated in brackets. Data pooled from two independent experiments, n=5-6 for each genotype in each experiment. Representative flow cytometry plots for CD4 vs CD44 **(C)**, IL-17 vs CD44 **(E)**, and IFNy vs CD44 **(G)** expression in CD4+ T cells following PMA/Ionomycin stimulation of blood samples collected on day 21 (EAE disease peak). Quantification of CD44+CD4+ **(D)**, CD44+IL-17+ **(F)**, and CD44+IFNy+ **(H)** cells as a frequency of CD4+ T cells in the blood. Absolute cell numbers for CD44+ IL-17+ **(I)**, CD44+ IFNy+ **(J)**, CD44+GM-CSF+ **(K)** and FOXP3+ **(L)** T cells in the CNS 28 days post-induction of EAE with MOG peptide for *Irf4*$^{+/+}$, *Irf4*$^{L116R/+}$, and *Irf4*$^{L116R/L116R}$ genotypes. Each dot is representative of a single mouse with symptomatic mice represented by triangles, n= 5-6 **(D-H)** or n=4-6 **(I-L)** per genotype. One-way ANOVA multiple comparisons are applied between groups. Logrank (Mantel-Cox) tests is applied for disease free survival curve comparison.

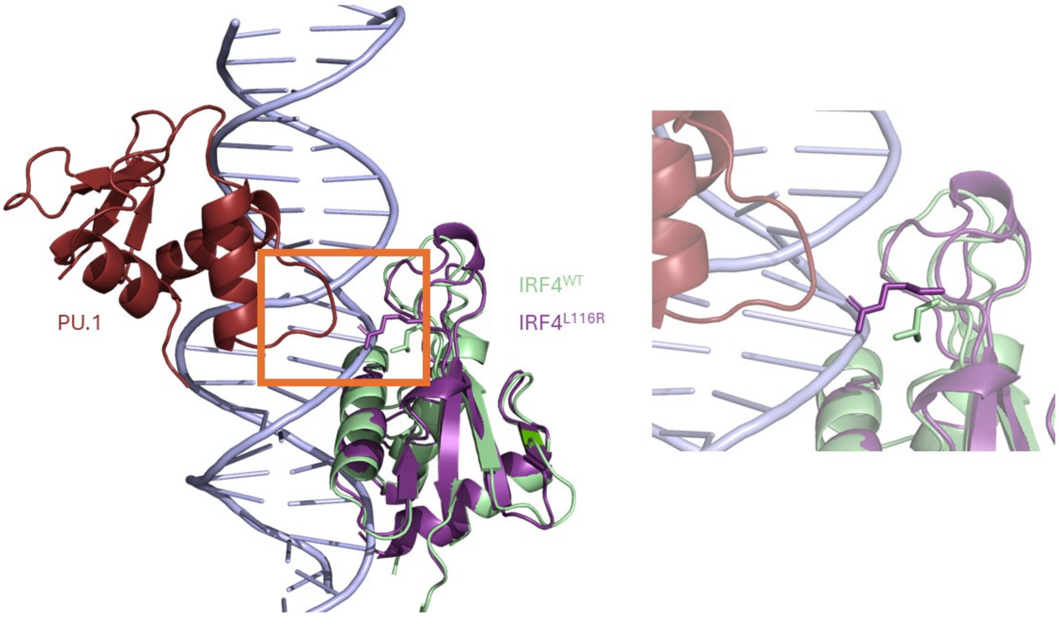


**Supplementary Figure 5. IRF4$^{L116R}$ modelled to alter the interaction of IRF4-DNA and potentially IRF4-PU.1. (A)** Structural representations of the IRF4 and PU.1 proteins in a complex with bound DNA with and without the L116R mutation. Images created using Pymol and PBD id: 7JM4.
